# Transient Zn^2+^ deficiency induces replication stress and compromises daughter cell proliferation

**DOI:** 10.1101/2023.12.08.570860

**Authors:** Samuel E. Holtzen, Elnaz Navid, Joseph D. Kainov, Amy E. Palmer

## Abstract

Cells must replicate their genome quickly and accurately, and they require metabolites and cofactors to do so. Ionic zinc (Zn^2+^) is an essential micronutrient that is required for hundreds of cellular processes, including DNA synthesis and adequate proliferation. Deficiency in this micronutrient impairs DNA synthesis and inhibits proliferation, but the mechanism is unknown. Using fluorescent reporters to track single cells via long-term live-cell imaging, we find that Zn^2+^ is required at the G1/S transition and during S-phase for timely completion of S-phase. A short pulse of Zn^2+^ deficiency impairs DNA synthesis and increases markers of replication stress. These markers of replication stress are reversed upon resupply of Zn^2+^. Finally, we find that if Zn^2+^ is removed during the mother cell’s S-phase, daughter cells enter a transient quiescent state, maintained by sustained expression of p21, which disappears upon reentry into the cell cycle. In summary, short pulses of mild Zn^2+^ deficiency in S-phase specifically induce replication stress, which causes downstream proliferation impairments in daughter cells.

**Significance:** Zinc is an essential micronutrient required for cells to grow and proliferate. However, the mechanism of how zinc influences proliferation is unknown. We show that short exposure to mild zinc deficiency in S-phase impairs DNA synthesis and induces replication stress, leading to pauses in daughter cell proliferation. However, pulses of low zinc during other phases of the cell cycle don’t affect mother cell cycle progression or daughter cell proliferation. These results indicate that while zinc is important for many proteins, during the cell cycle short pulses of mild zinc deficiency have the biggest impact on a cell’s ability to synthesize DNA, suggesting that DNA polymerase complex acts as a gate keeper, sensing zinc status in the cell.

## Introduction

The mammalian cell cycle is composed of several mostly irreversible ratchet-like steps that prepare a cell for division (1–3). Mother cells must grow and license origins of replication in G1, replicate their genome efficiently and accurately in S-phase, and ensure that each genome is duplicated once and only once in G2 before dividing into two daughter cells. In response to stress, loss of adhesion, DNA damage, or mitogen withdrawal, cells may elongate any one of these phases or temporarily leave the cell cycle and enter a quiescent state after division, termed G0 (4–7). Cells constantly evaluate their intra- and extracellular state, including metabolite, nutrient and energy balance, and integrate these signals to make go/no-go decisions at the transition between each of these phases (5, 8, 9).

Ionic zinc (Zn^2+^) is required for human health (10). Ten percent of human genes are predicted to encode proteins that have one or more Zn^2+^ binding sites, where Zn^2+^ is suggested to play a role in structure stabilization, catalysis, or signaling (11). Recently, we established that Zn^2+^ is an important and enigmatic signaling ion in regulation of cellular processes. For example, cells transiently increase their cytosolic labile Zn^2+^ in early G1 and require optimal labile Zn^2+^ for proliferation (12). Further, elevation of cytosolic Zn^2+^ activates flux through the mitogen-activated protein kinase (MAPK) pathway (13). Finally, Zn^2+^ induces changes in transcription in mouse hippocampal neurons (14, 15). At a cellular level, deficiency of this essential micronutrient can cause a loose constellation of varied symptoms such as DNA synthesis impairment, oxidative stress, and apoptosis (16–19). Several studies have attempted to dissect how Zn^2+^ deficiency can cause its hallmark effects in humans through the lenses of DNA synthesis, cell cycle protein expression and the DNA damage response (16, 20, 21). all of these studies, researchers found that Zn^2+^ deficient cells are unable to proliferate.

Early studies identified the requirement of Zn^2+^ for DNA synthesis and proliferation in several cell types (16, 20), and determined that treating cells with an extracellular Zn^2+^ chelator inhibits proliferation through inhibition of cyclin mRNA expression (21). More recent work has determined that MCF10a cells experiencing mild Zn^2+^ deficiency bifurcate into two populations after cell division depending on how long they were Zn^2+^ deficient before dividing (22). In the first population, cells enter quiescence and are able to enter the cell cycle again upon resupply of Zn^2+^. This population seems to increase in abundance with the duration of Zn^2+^ deficiency. In the second population, cells are able to pass the restriction point, only to stall in a proliferative state characterized by hyperphosphorylation of the Rb protein, indicating these cells are indeed committed to another cell cycle but unable to complete it.

These studies pose several unanswered questions. Since cells bifurcate into two populations after division depending on how long cells were exposed to Zn^2+^ deficiency, is Zn^2+^ required at specific phases of the cell cycle, or do cells integrate time spent in Zn^2+^ deficiency to determine cell fate? What causes Zn^2+^ deficient cells to accrue DNA damage and show decreased DNA synthesis? Finally, what molecular processes contribute to the daughter cell proliferation/quiescence decision? Using a combination of fluorescent reporters for single cell tracking, long term live-cell imaging, immunofluorescence, and flow cytometry, we show that Zn^2+^ is required at the G1/S transition and in S-phase for timely S-phase completion. We then show that a short pulse of mild Zn^2+^ deficiency reversibly inhibits DNA synthesis, leads to accumulation of replication protein A subunit 2 (RPA2) foci, and activates the S-phase checkpoint. Analysis of likely causes of replication stress implicates an essential role for Zn^2+^ in the function of DNA polymerase. Finally, we show that Zn^2+^ deficiency in the mother cell’s S-phase induces temporary daughter cell quiescence through sustained expression of p21. Together, our results indicate that acute, mild, and temporary Zn^2+^ deficiency in S-phase, but not other cell cycle phases, causes reversible replication stress, which leads to daughter cell entry into quiescence after division.

## Results

### Zn^2+^ is required at the G1/S transition and in S-phase for normal S-phase progression

Zn^2+^ deficient cells show a bifurcation in cell fate after cell division, shifting their fate away from proliferation and towards quiescence with increased duration of Zn^2+^ withdrawal (22). To simulate mild Zn^2+^ deficiency, previous work used the membrane permeable Zn^2+^ chelator tris(2-pyridylmethyl)amine (TPA) in culture media, which lowers the labile Zn^2+^ levels in MCF10a cells from 80 pM to 1 pM (12, 22). In this study, two factors confound interpretation of this prior work: the timing and the duration of the treatment. To address this, we used a pulse-chase treatment strategy coupled with the FUCCI (CA) sensor, which is able to resolve cell cycle phase boundaries and therefore allow accurate measurement of cell cycle phase lengths (23). The sensor is composed of two fluorescent proteins fused to fragments of human Cdt1 and Geminin proteins, which are reciprocally ubiquitinated and degraded in a cell cycle phase specific manner (Figure 1A) (23). Using long-term live-cell imaging, we tracked MCF10a cells through several cell divisions using a published pipeline (24), extracted single-cell fluorescence intensities of the two probes, and identified the cell cycle phase at each frame (Figure 1B).

**Figure 1:**
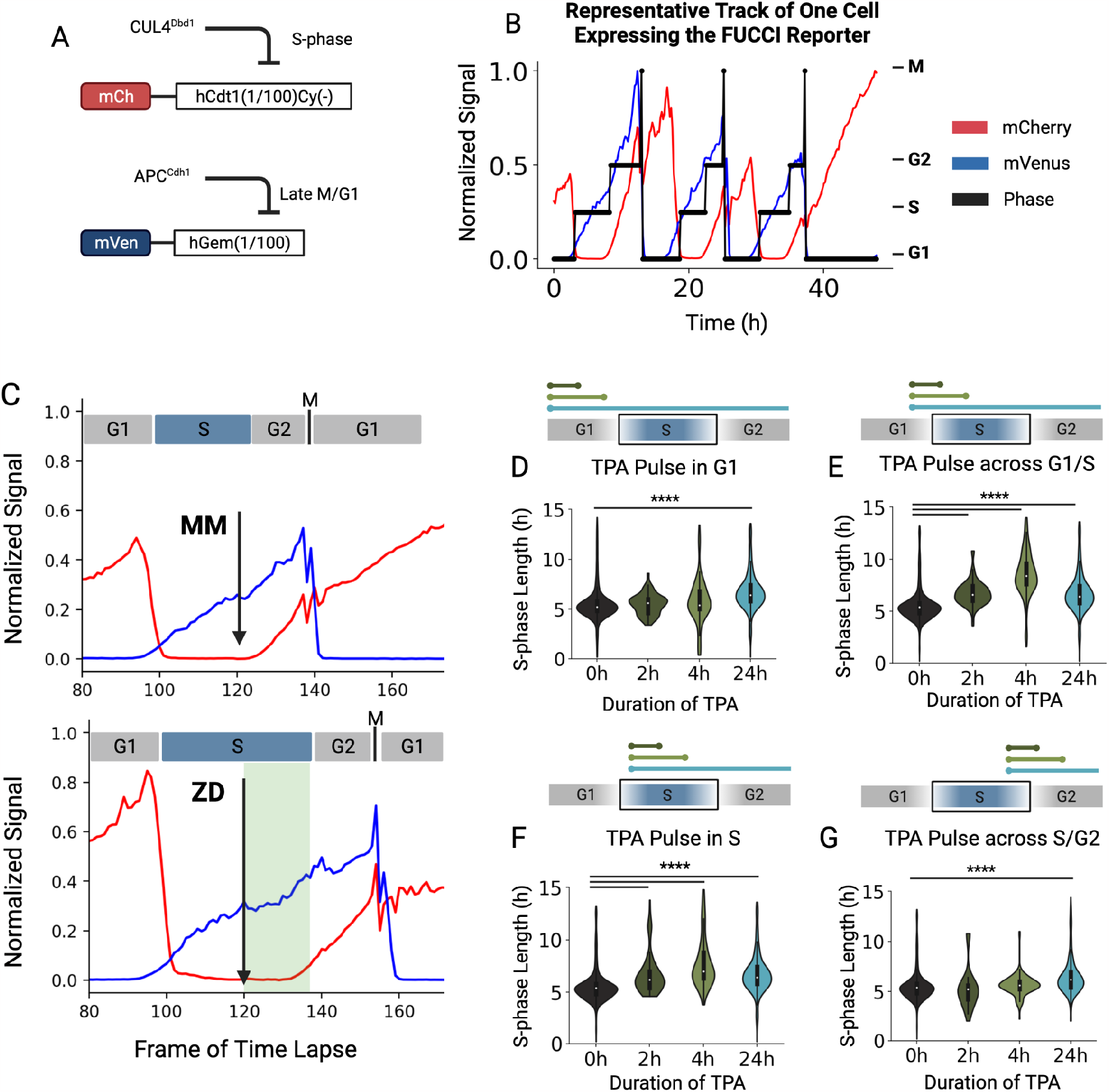
Cells treated with Zn^2+^ deficient media across the G1/S transition and during S-phase have an elongated S-phase. (A) A schematic of the FUCCI (CA) sensor mechanism of operation. The mCherry and mVenus fluorescent proteins are conjugated to fragments of the human Cdt1 and human Geminin proteins, which are reciprocally targeted for degradation by the CUL4^Dbd1^ and APC^Cdh1^ ubiquitin ligases during S-phase and G1, respectively. (B) A representative track of a cell expressing the FUCCI (CA) sensor throughout three cell cycles. The mCherry and mVenus intensities are min-max normalized. The cell cycle phases assigned *in silico* are shown as the black stair-step lines. (C) Representative traces of cells treated with MM media (top) or ZD media (bottom) during S-phase. Shaded area represents the duration and timing of ZD exposure. (D) S-phase lengths of cells exposed to ZD media during G1, and whose Zn^2+^ was resupplied in G1. (E) S-phase lengths of cells exposed to ZD media across the G1/S transition. (F) S-phase lengths of cells exposed to ZD during S-phase, and whose Zn^2+^ was resupplied in S-phase. (G) S-phase lengths of cells exposed to ZD across the S/G2 transition. **** = p<0.001. n=3 biological replicates. Colored bars above the cell cycle phase diagrams indicate the timing and duration of the Zn^2+^ deficiency pulse.

We pulsed MCF10a cells expressing the FUCCI (CA) sensor with Zn^2+^ deficient media containing 3μM TPA for 2 or 4 hours, then chased with Zn^2+^ adequate media. After cell tracking and signal extraction, we were able to identify changes in cell cycle phase lengths as a function of Zn^2+^ deficiency duration. If Zn^2+^ was withdrawn from cells at the G1/S transition and in S-phase, cells had a significantly longer S-phase than untreated cells (Figure 1C, E-F). This effect increased with the duration of Zn^2+^ withdrawal. If Zn^2+^ was withdrawn and replaced while cells were still in G1, there was no effect on the subsequent S-phase, indicating that Zn^2+^ is not required in G1 for timely S-phase completion (Figure 1D). Similarly, if Zn^2+^ was withdrawn in S-phase and resupplied in G2, S-phase was not elongated, implying that Zn^2+^ is not required in late S-phase for timely S-phase progression (Figure 1G). Together, these data illustrate a specific Zn^2+^ requirement at the G1/S transition and in S-phase for timely S-phase progression, and that a process at this stage is disrupted when deprived of adequate Zn^2+^.

### Zn^2+^ deficient cells show impaired DNA synthesis

After 24 hours of Zn^2+^ deficiency, proliferative daughter cells show strongly impaired DNA synthesis rates compared to Zn^2+^ replete cells (22). Other studies have shown that Zn^2+^ is required for DNA synthesis in a number of cell types (16, 20). Since we saw an impaired S-phase in mother cells exposed to pulses of Zn^2+^ deficiency, we wanted to determine whether this elongated S-phase is coupled with inhibited DNA replication. We therefore used a pulse of 5-ethynyl-2′-deoxyuridine (EdU) coupled with click-chemistry mediated addition of a fluorescent dye and flow cytometry to measure DNA synthesis rate of single cells (25). In brief, cells were incubated with the Zn^2+^ chelator TPA for two or four hours, then exposed to EdU to measure incorporation of EdU within a defined time window. After fixation and counterstaining with DAPI for DNA content, we identified the DNA synthesis rate in single cells (25). We found that two hours of Zn^2+^ deficiency inhibits DNA synthesis by half, and that this inhibition is also observed during a four-hour pulse of Zn^2+^ deficiency (Figure 2A-B). Resupply of Zn^2+^ fully reverses the DNA synthesis inhibition, demonstrating this DNA synthesis defect is reversible (Figure 2B).

**Figure 2:**
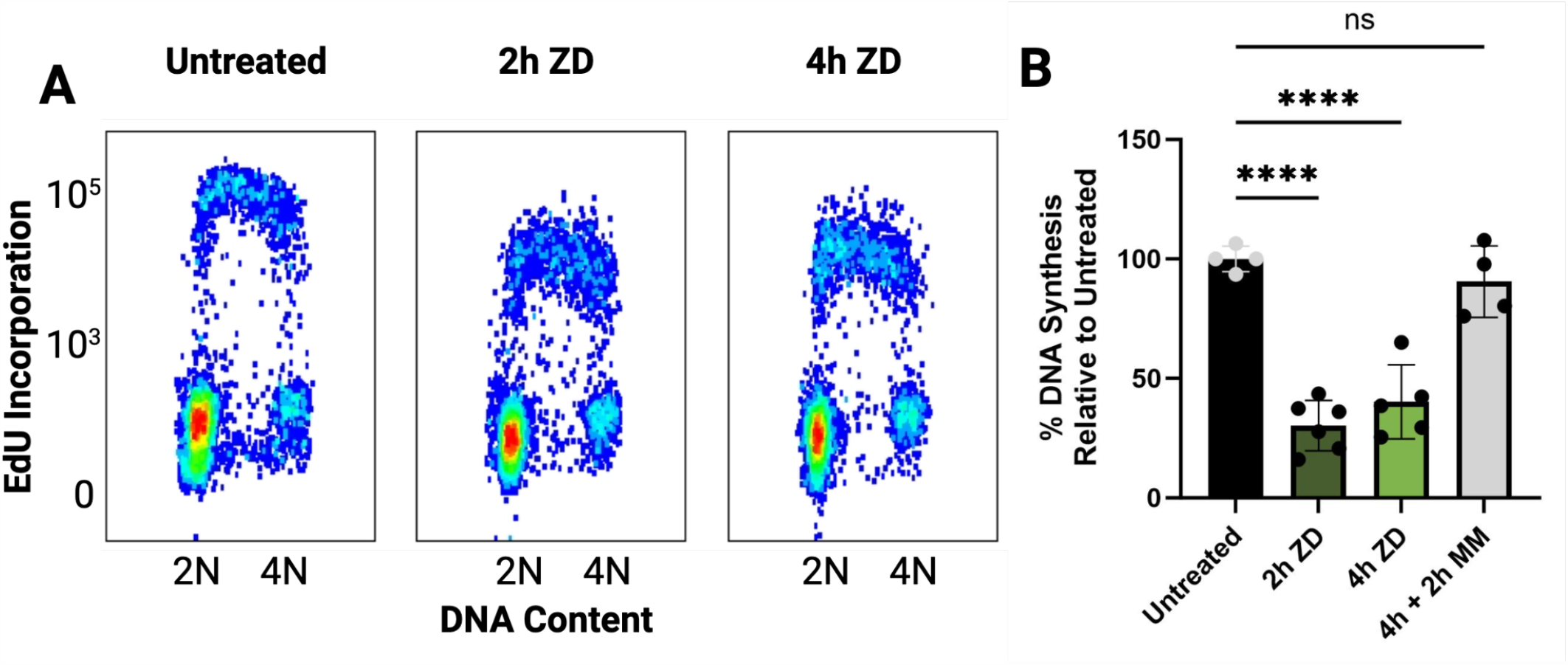
Zn^2+^ deficient cells show impaired DNA synthesis. (A) Flow cytometry plots showing Zn^2+^ adequate and Zn^2+^ deficient cells treated with the nucleotide analog EdU to probe DNA synthesis rate, then counterstained with the DNA content dye DAPI. Each point represents a single cell. (B) Quantification of the plots in A. Each point is the median EdU signal of S-phase cells (EdU+) in one replicate. All data was then normalized to the untreated condition. Error bars represent the 95% confidence interval. *** = p < 0.005, **** = p < 0.001. n ≥ 4 biological replicates.

### Zn^2+^ deficient cells show signs of replication stress

Impaired DNA synthesis during S-phase is termed replication stress, and can be due to several disparate factors such as DNA adducts (26, 27), nucleotide depletion (9, 28), or inhibition of the DNA polymerase (29, 30). Inhibition of the DNA polymerase and DNA lesions can lead to uncoupling of the Cdc45, Mcm2– 7, GINS (CMG) helicase from DNA polymerase (31, 32). This leads to long tracts of single-stranded DNA, which are rapidly coated by RPA2 (Figure 3A). In order for functional uncoupling between the CMG helicase and DNA polymerase to occur, the helicase must be active and the polymerase must be stalled (31, 32). To determine whether the CMG helicase is still active, but the polymerase is stalled by low Zn^2+^, we measured whether RPA2 foci accumulate, which indicates single-stranded DNA accumulation. To do this, we pulsed cells expressing the FUCCI (CA) sensor with Zn^2+^ deficient media, then fixed and stained them for RPA2. To identify the cell cycle phases of each fixed cell, we used a Gaussian mixture model (GMM) with the *z*-score normalized mVenus and mCherry signal in each cell, then assigned each cell cycle phase by hand (Figure S1).

**Figure 3:**
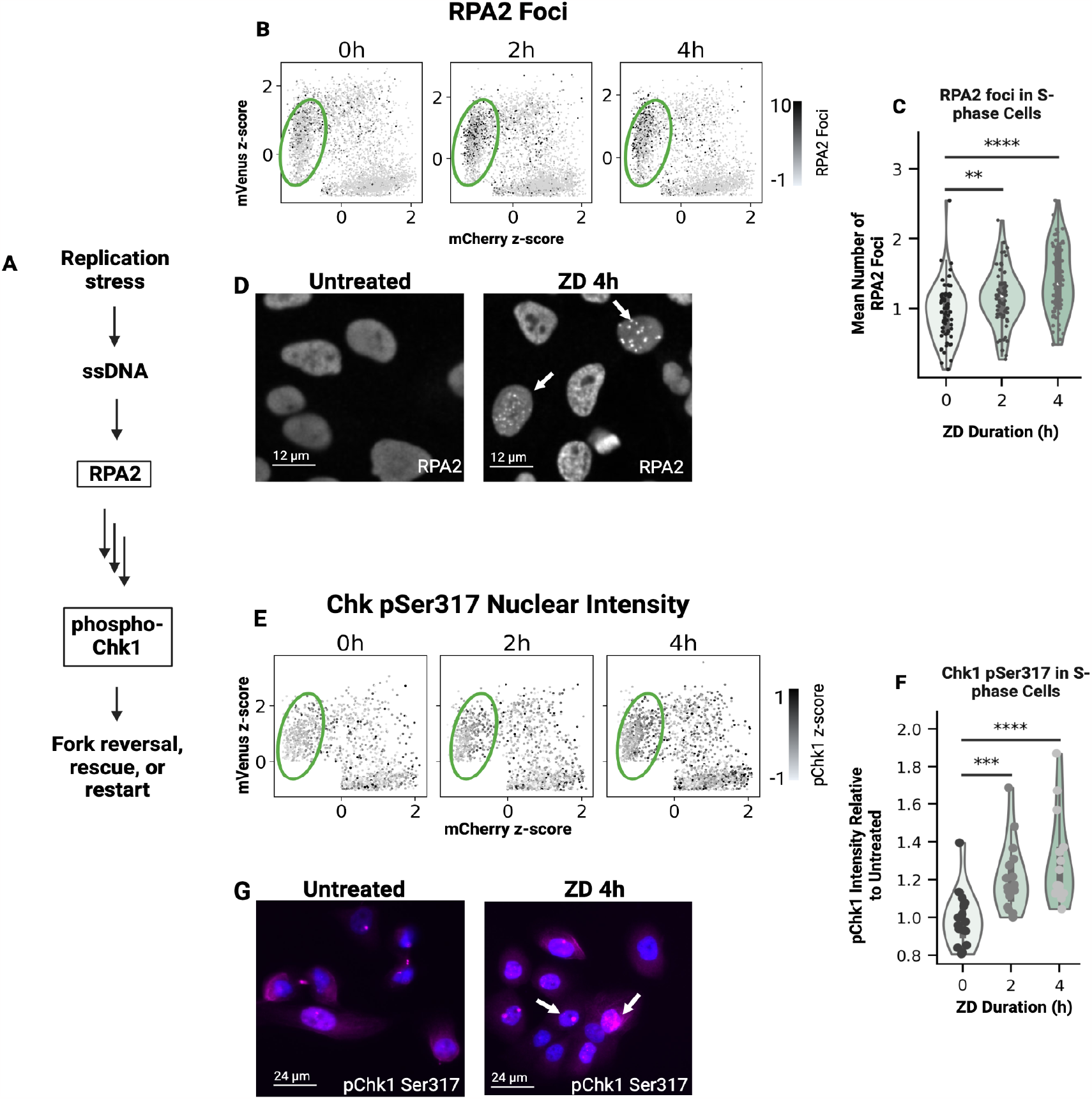
Acute Zn^2+^ deficiency increases replication stress markers. (A) Pathway of sensing and response to replication stress. Replication stress causes accumulation of single-stranded DNA (ssDNA), which is coated by RPA2. Downstream proteins sense this accumulation and signal activation of Chk1 via phosphorylation, which activates effector proteins to stabilize and restart replication forks. (B) Quantitative imaging-based cytometry plots of mean nuclear fluorescence of the mCherry and mVenus construct of the FUCCI (CA) sensor on the horizontal and vertical axes, respectively. Each point is one cell, and the gray level of each point corresponds to the number of RPA2 foci it has. Green ovals indicate cells in S-phase. (C) Quantification of RPA2 foci in S-phase cells after treatment with Zn^2+^ deficient media. Each point is the average number of RPA2 foci in S-phase cells in one well of a 96-well plate. (D) Representative images of RPA2 foci in untreated cells and cells treated with Zn^2+^ deficient media for 4h. White arrows indicate cells with RPA2 foci. (E) Quantitative imaging-based cytometry plots of mean nuclear fluorescence of the mCherry and mVenus construct of the FUCCI (CA) sensor on the horizontal and vertical axes, respectively. Each point is one cell, and the gray level of each point corresponds to its mean nuclear phospho-Chk1 (Ser317) intensity. Green ovals indicate cells in S-phase. (F) Quantification of pChk1 signal in S-phase cells after treatment with Zn^2+^ deficient media. Each point is the average phospho-Chk1 (Ser317) signal in S-phase cells in one well of a 96-well plate. (G) Representative images of phospho-Chk1 (Ser317) signal in untreated cells and cells treated with Zn^2+^ deficient media for 4h. Blue indicates nuclear DAPI staining. Magenta indicates phospho-Chk1 Ser317 signal. White arrows indicate cells with phospho-Chk1 (Ser317) signal. ** = p < 0.01, *** = p < 0.005, **** = p < 0.001. n ≥ 2 biological replicates.

Exposure of cells to a 2 or 4 hr pulse of Zn^2+^ deficiency induced an increase in the number of RPA2 foci in S-phase cells, which increased with the duration of treatment (Figure 3B-D). RPA2 foci, much like DNA synthesis impairment, were resolved with resupply of Zn^2+^ (Figure S2). Because RPA2-bound singlestranded DNA leads to activation of the ATR-Chk1-p53 signaling pathway (31), we probed phosphorylation of the Chk1 kinase at Ser317 by immunofluorescence. We found that a 2 hr or 4 hr pulse of Zn^2+^ deficiency increased Chk1 phosphorylation in S-phase along with RPA2 foci accumulation and DNA synthesis impairment (Figure 3E-G). These results indicate that RPA2 accumulation is accompanied by increased signaling through the Chk1 pathway in response to transient Zn^2+^ deficiency in S-phase.

### Zn^2+^ deficiency does not cause double-stranded breaks, origin under-licensing, replication fork slowing, or an imbalance in the nucleotide pool

Since replication stress can be due to replication origin under licensing in G1 (33), it is possible that Zn^2+^ deficiency impaired licensing, which in turn could contribute to the accumulation of RPA2 foci. We tested whether Zn^2+^ deficiency causes under licensing of origins using a flow cytometry assay that probes for chromatin bound MCM2 as described previously (33). Transient Zn^2+^ deficiency did not cause under licensing of replication origins, further supporting that the MCM2 helicase is loaded at replication origins and is functional and that Zn^2+^ deficiency is not interfering with S-phase preparation (Figure S3).

ATR-Chk1 activation can decrease rates of DNA synthesis in response to replication stress (32). To determine whether the ATR-Chk1 signaling axis globally inhibited DNA replication under conditions of Zn^2+^ deficiency, we co-treated cells with Zn^2+^ deficient media and the Chk1 inhibitor CHIR-124 for 4 hrs. After calculating the median EdU incorporation of each sample compared to the untreated controls, we found that, although DNA synthesis rate was slightly higher in cells treated with CHIR-124, it was not a full rescue as was the case with Zn^2+^ resupply (Figure S3E, Figure 2B). Therefore, Chk1 activation seems to be a consequence of Zn^2+^ deficiency-induced DNA synthesis impairment, not a cause of it.

Double-stranded breaks (DSBs) are genotoxic lesions that activate the DSB repair pathway mediated by the ATM/Chk2 signaling axis and repair the damage using non-homologous end joining or homologous recombination. DSBs in S-phase can induce replication stress through the ATM/Chk2 signaling axis, or through co-activation of Chk1 and Chk2 (6, 32, 34). To determine whether acute Zn^2+^ deficiency induces double-stranded breaks, we treated FUCCI(CA) expressing cells with Zn^2+^ deficient media and probed for 53BP1, a central player in the double-stranded break response, in fixed cells using immunofluorescence. We found that four-hour Zn^2+^ deficiency does not increase 53BP1 foci relative to the untreated control in any cell cycle phase (Figure S3A). Further, we determined that 24-hour Zn^2+^ deficiency does not activate the checkpoint kinase Chk2 relative to the untreated control, indicating that Zn^2+^ deficiency does not exert its effects through double-stranded break repair signaling, but rather through replication stress signaling (Figure S4B).

An imbalance in the intracellular nucleotide pool can activate the replication stress pathway via Chk1 (28, 28). To determine if acute Zn^2+^ deficiency changes the nucleotide pool, we conducted untargeted metabolomics of polar metabolites. We found that, although there were small changes in nucleotide precursors, there was no change in the concentrations of free nucleotides, nor in their deoxy counterparts (Figure S5). This indicates that Zn^2+^ deficiency does not impair DNA synthesis through depletion or imbalance of the nucleotide pool.

### Transient Zn^2+^ deficiency during S-phase of the mother cell impairs proliferation of daughter cells

Cells integrate internal and external signals during the mother cell cycle in order to dictate daughter cell proliferation after division (5, 35). Since endogenous replication stress in the mother cell can inform daughter cell proliferation (4), we sought to identify the consequences of Zn^2+^-deficiency-induced mother cell replication stress on daughter cell proliferation. To do this, we used an analysis technique presented in Min et al (5). This technique takes advantage of a CDK2 activity sensor’s ability to resolve cell cycle commitment (Figure 4A) (36). After division, cells take on three fates: In the first, cells are born with a high CDK2 activity and hyper-pRb, which indicates that they are born committed to another cell cycle (Figure 4A, CDK2_inc_, blue). In the second, cells are born with a low CDK2 activity, but cross the 0.5 arbitrary unit (AU) threshold and commit to another cell cycle sometime later (Figure 4A, CDK2_emerge_, yellow). In the third, cells are born with a CDK2 activity below 0.5 and remain there for the rest of the time-lapse, indicating they are in a quiescent state that is not reversed during the time-lapse (Figure 4A, CDK2_low_, red). Cell traces from asynchronously cycling cells can be aligned to mitosis such that the three populations in each mitosis “slice” can be identified and expressed as a fraction. This is repeated for all frames of the time lapse to yield a map of when TPA acts to influence cell fate after division (Figure 4B).

**Figure 4:**
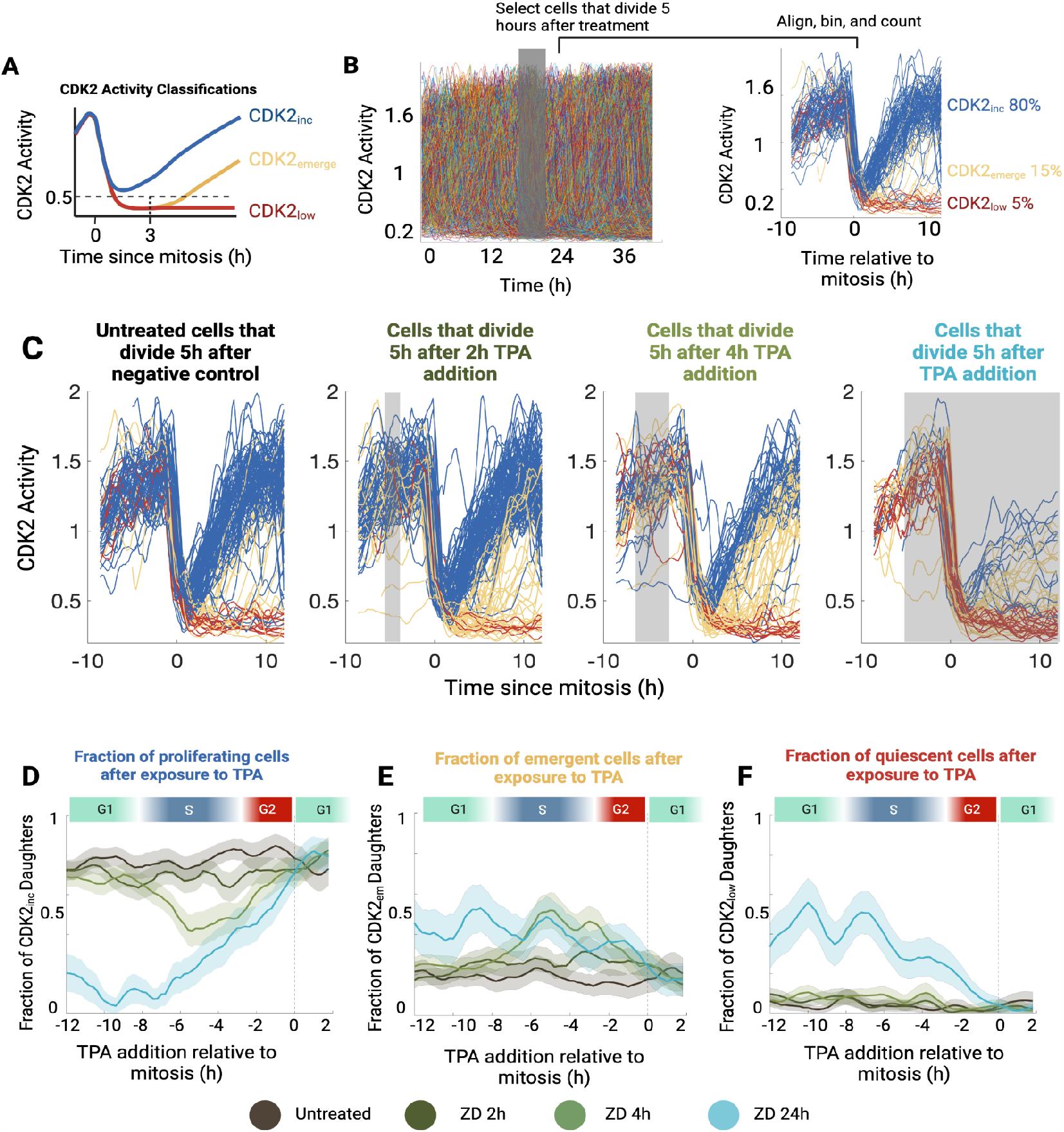
Zn^2+^ deficiency in the mother cell S-phase causes proliferation pause in daughter cells. (A) Schematic of the CDK2 activity sensor classifications. CDK2_inc_ indicates a CDK2 activity that is higher than 0.5 after mitosis. CDK2_emerge_ indicates a brief pause in G0, then entry into G1. CDK2_low_ indicates an induction into quiescence that persists for the rest of the movie. (B) Analysis scheme of the CDK2 activity pulse-chase experiment. Cells are binned according to the time they divide relative to their treatment with TPA. Cells are then binned according to their CDK2 activity after division. Gray shaded area indicates treatment timing and duration. (C) Single-cell traces of cells expressing the CDK2 sensor aligned to mitosis. All cells shown were treated with Zn^2+^ deficient media 5 hours before mitosis. The duration of this pulse is indicated by the gray rectangle in each plot. Cells are color-coded based on their CDK2 activity 15 frames after anaphase. Each plot contains 100 single cell traces. (D) Plot of fraction of proliferating daughter cells as a function of mother cell age at treatment addition. Dashed line indicates mitosis. Light colored bands represent 95% confidence intervals. (E) Plot of fraction of CDK2_emerge_ daughter cells as a function of mother cell age at treatment addition. Dashed line indicates mitosis. Light colored bands represent 95% confidence intervals. (F) Plot of fraction of CDK2_low_ daughter cells as a function of mother cell age at treatment addition. Dashed line indicates mitosis. Light colored bands represent 95% confidence intervals. n=2 biological replicates. Each time point contains the average and 95% CI of at least 20 individual cell traces.

To illustrate this process, we selected cells that divided five hours after TPA addition from each treatment group and identified the distribution of cell fates after division. As shown previously (4, 5, 37), we found that 80% of untreated MCF10a cells increase their CDK2 activity immediately after mitosis, indicating that they are born committed to the cell cycle (Figure 4C). The other 20% enter a brief resting state, deemed CDK2_emerge_. A pulse of 2 or 4 hours of Zn^2+^ deficient media five hours before cell division leads to progressive loss of the CDK2_inc_ population (blue), and a concomitant increase in the proportion of CDK2_emerge_ cells (Figure 4C). Finally, if Zn^2+^ deficient media was added during S-phase and kept until the end of the time lapse, cells bifurcate into two populations, where the majority either enter the CDK2_emerge_ or CDK2_low_ cell fate (Figure 4C). This agrees with previous research (22).

After repeating this analysis for all frames of the time lapse, we found that 80% of MCF10a cells are born proliferative, which agrees with prior research (4, 5, 37). In contrast, cells that are pulsed with the Zn^2+^ chelator TPA during S-phase (five hours before mitosis) for two or four hours show a decrease in proliferative daughter cells (Figure 4C). This decrease is accompanied by a proportional increase in the fraction of CDK2_emerge_ daughter cells, indicating that if cells lack Zn^2+^ in S-phase, they enter a transient state of low CDK2 activity after cell division (Figure 4E). Critically, these cells do not enter a permanent CDK2_low_ state after division, indicating that they are not quiescent to the end of the time lapse, but are instead in a transient state of cell cycle exit (Figure 4F). Importantly, beginning the pulse of Zn^2+^ deficiency after the S/G2 boundary or before mid S-phase has no effect on the proliferation of daughter cells. In contrast, the cells that are treated with TPA until the end of the time lapse show a strong propensity to be either CDK2_emerge_ or CDK2_low_ after division, as shown previously (Fig 4E-F) (22). The distribution of fates shifts toward the CDK2_low_ population the longer a cell is in Zn^2+^ deficient media (Figure 4E-F). Of note, the transmission of replication stress through mitosis is not accompanied by mitotic DNA synthesis (MiDAS) (Figure S7), which marks the transmission of unresolved replication stress to daughter cell quiescence (38, 39). These data demonstrate that cells pulsed with TPA during S-phase show a steady decrease in proliferative cells after division and reach a nadir during the middle of S-phase. This impairment is sustained during the latter half of S-phase until the S/G2 transition. Thus, cells that are Zn^2+^ deficient in the latter half of S-phase enter a transient CDK2_emerge_ state after division, implying that the replication stress and DNA synthesis inhibition due to low Zn^2+^ conditions are directly impacting daughter cell proliferation.

### Sustained p21 expression maintains cell cycle pause induced by Zn^2+^ deficiency

Transient Zn^2+^ deficiency in the mother cell S-phase causes a proliferative pause in the daughter cells (Figure 4). The tumor suppressor p21, among other such proteins, acts as a competitive inhibitor of the active Cyclin E-CDK2 complex (1), and thus is one factor that can prevent cells from entering the cell cycle. p21 acts to maintain the quiescent state in both unperturbed cells and cells who inherit damaged DNA from mother cells (6, 35, 37). If cells are pulsed with low Zn^2+^ in G1 of the mother cell (approximately 12 hours before mitosis), daughter cell proliferation is not significantly different from the daughters of Zn^2+^ adequate cells (Figure 5A). If cells are pulsed with low Zn^2+^ in S-phase (approximately 6 hours before mitosis), and not resupplied prior to the end of S-phase (at approximately 2 hours prior to mitosis), the fraction of daughters born proliferative is reduced to approximately 45% (Figure 5A). We therefore hypothesized that transient Zn^2+^ deficiency in S-phase would induce a sustained p21 expression after division when compared to the untreated cells. We similarly hypothesized that transient Zn^2+^ deficiency during G1 would induce p21 expression dynamics similar to the untreated cells, since we do not see impaired proliferation. To test this, we used a cell line expressing the CDK2 activity sensor, H2B-mTurquoise2 as a nuclear marker, and a p21-mCitrine construct expressed at the endogenous locus (Figure 5B) (37). We treated these cells with Zn^2+^ deficient media for four hours, followed by a resupply of Zn^2+^ adequate media. We then analyzed cells that divided 12 or 6 hours after transient Zn^2+^ deficiency for changes in p21 dynamics in both CDK2_inc_ and CDK2_emerge_ populations.

**Figure 5:**
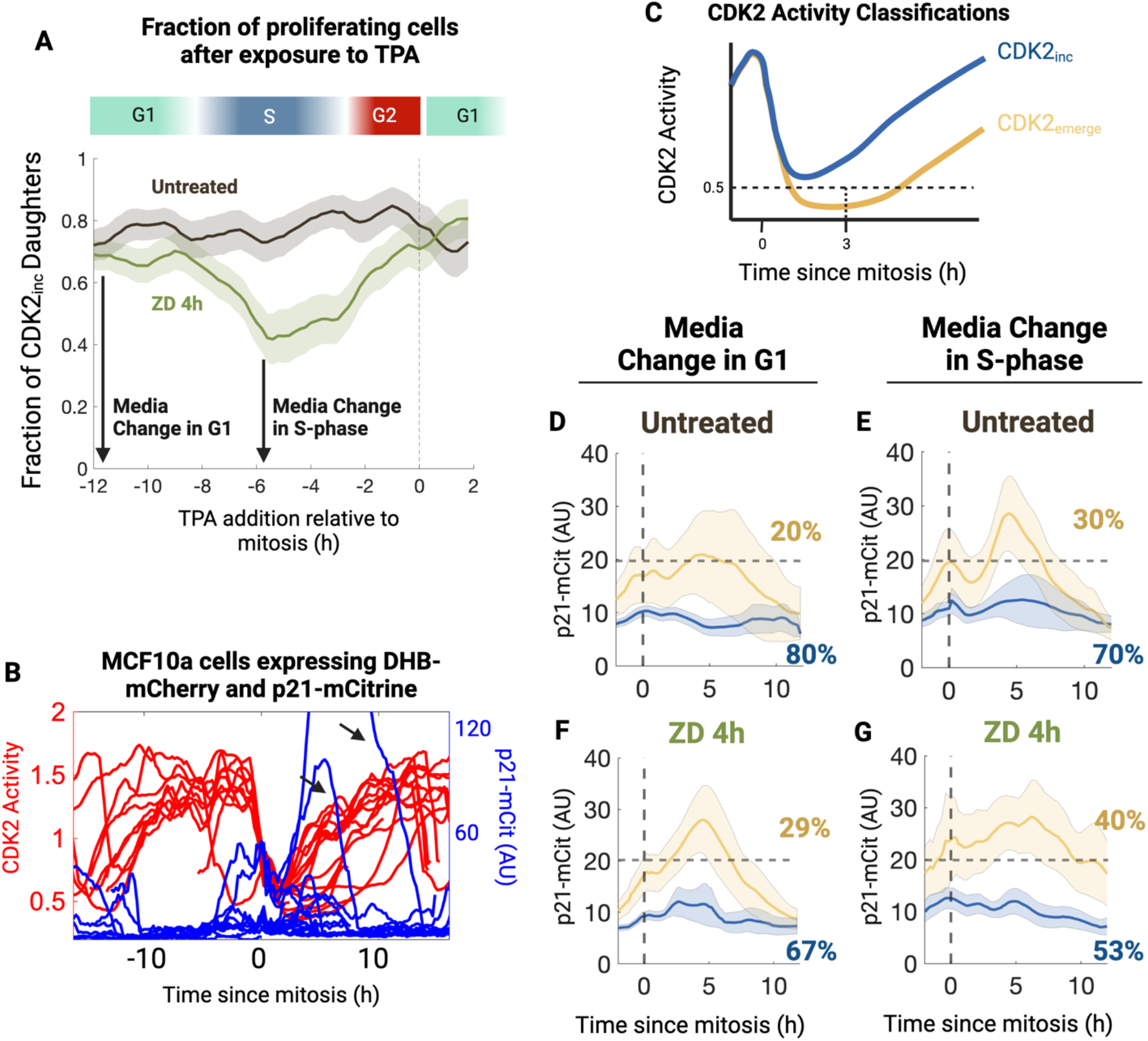
Transient Zn^2+^ deficiency during S-phase induces proliferation impairment accompanied by sustained p21 expression in G0. (A) Results from Figure 5D. Arrows indicate times relative to anaphase that induces normal daughter cell proliferation and impaired daughter cell proliferation, respectively. (B) Representative traces of cells expressing DHB-mCherry (red) and p21-mCitrine (blue) from the endogenous locus aligned to anaphase. n=10 single-cell traces. Eight traces are born with low p21 expression. Two traces (black arrows) show transiently increased p21 expression. (C) CDK2 activity classification scheme. Yellow traces indicate CDK2_emerge_ population, blue traces indicate CDK2_inc_ population. (D, F) Average p21 expression in cells that divided 60 frames (∼ 12 hours) after treatment with either a 4 hr pulse of Zn^2+^ deficient media or control media, color coded to show the CDK2_inc_ (blue) and CDK2_emerge_ (yellow) populations. Light bands indicate 95% confidence intervals. n = 2 biological replicates. (E, G) Average p21 expression in cells that divided 5 hours after treatment with a 4-hour pulse of Zn^2+^ deficient media or control media, color coded to show the CDK2_inc_ (blue) and CDK2_emerge_ (yellow) populations. Light bands indicate 95% confidence intervals. n = 2 biological replicates. Vertical dashed lines represent the frame of mitosis. Horizontal dashed lines indicate a 20 AU threshold to highlight increased and sustained p21 dynamics.

Eighty percent of untreated cells are born as CDK2_inc_ with low p21 expression (blue-colored traces) (Figure 5B,D,E), which also agrees with previously published work (36, 37). The other 20% of cells are born as CDK2_emerge_ and show a transient increase in p21 that falls to basal expression after seven hours, followed by an increase in CDK2 activity and commitment to another cell cycle (Figure 5B,D). This also agrees with previously published results showing that in asynchronously cycling cells, a small population of cells accumulates DNA damage and pauses the cell cycle via expression of p21 (37). Similar to the untreated condition, when Zn^2+^ was depleted and resupplied in G1 (12 hours prior to mitosis), the CDK2_emerge_ population shows comparable p21 dynamics to CDK2_emerge_ cells in the untreated condition (Figure 5F). Although there are fewer proliferative cells in this population (Figure 5F), the p21 signal rises and returns to baseline with dynamics similar to the untreated cells. Therefore, a pulse of Zn^2+^ deficiency in G1 shows similar p21 dynamics to untreated cells. When Zn^2+^ is depleted and resupplied during S-phase, however, the fraction of cells born CDK2_emerge_ increases to approximately 40%, which agrees with the data in Figure 4D. Further, this population shows sustained p21 expression that does not drop to basal levels for at least twelve hours after division (Figure 5E,G), indicating that the CDK2_emerge_ state is being maintained by sustained p21 expression. This result indicates that Zn^2+^ deficiency induced replication stress during S-phase of the mother cell is inherited by daughter cells to cause temporary quiescence.

## Discussion

Duplicating a genome is a daunting task and requires careful monitoring of cell state (40). Human cells constantly sample their intra- and extracellular environment for growth factors, genotoxic stress, and metabolites and integrate these signals to determine whether to repair, grow, or die (5, 9, 35). Early work to identify the role of Zn^2+^ deficiency in the mammalian cell cycle revealed a loosely-related set of phenotypes ranging from DNA synthesis impairment (21) to oxidative stress (18, 19) to apoptosis (17). None of these studies simulated mild, physiologically relevant Zn^2+^ deficiency, and instead relied on high concentrations of metal chelators and cell synchronization, which are known to disrupt cell cycle progression via stress responses (16, 20, 21, 33, 41). Due to zinc’s role in a multitude of cellular processes, it is often assumed that Zn^2+^ deficiency will induce pleiotropic effects on cells. To dissect whether Zn^2+^ acts during a specific cell cycle phase to impair proliferation, we used pulse-chase treatments along with fluorescent cell cycle reporters. We found that transient Zn^2+^ deficiency impaired S-phase progression, slowed DNA synthesis rates, and induced replication stress signaling, which contributed to a pause in the cell cycle of daughter cells. Importantly, transient Zn^2+^ deficiency during other phases of the cell cycle had no adverse effects. These results demonstrate a specific requirement for Zn^2+^ during S-phase, in contrast to the assumption that Zn^2+^ deficiency affects cellular processes pleiotropically.

Replication stress can be caused by DNA lesions and adducts (32), imbalanced nucleotides (9, 28), underlicensed replication origins (33, 39, 40), inhibition of DNA polymerase (31), or endogenous stochastic uncoupling of replication forks (4). The results of many previous studies into Zn^2+^ and in the mammalian cell cycle revealed phenotypes that could be classified as “replication stress,” including impaired DNA synthesis, increased DNA damage signaling and production of oxidative products (18, 20, 21). In this study, we found that when cells were treated with a short pulse of Zn^2+^ deficiency, there was no evidence of under licensing, nucleotide imbalance, or double-stranded breaks. Instead, we found that acute Zn^2+^ deficiency increases RPA2 recruitment to nuclear foci and activates the checkpoint kinase Chk1. RPA2 recruitment and Chk1 activation are both hallmarks of functionally uncoupled polymerase-helicase complexes (31, 32, 42), indicating that the helicase is still fully functional under conditions of Zn^2+^ deficiency.

The DNA polymerases Polδ, Polδ, and Polε are protein complexes in the eukaryotic replisome responsible for synthesizing the genome, and the catalytic subunits of these complexes have at least one Zn^2+^ binding motif that appears to be critical for their function (43). For example, the zinc finger domain of the *S. cerevisiae* Polε is important for transducing DNA damage signaling through the S/M checkpoint pathway, and seems to impair signaling when mutated (44). Further, the C-terminal zinc finger of the *S. cerevisiae* Polδ is required for its association between other subunits of the Polδ complex, and mutation of the zinc finger abolishes activity *in vivo* (45). Finally, human Polδ contains a C-terminal zinc finger, and treatment with cisplatin displaces the Zn^2+^ ion and inhibits its activity *in vitro* (46). In addition to the role of zinc fingers within the Pol subunits, there is evidence that suggests Zn^2+^ plays a role in stabilizing intermolecular interactions formed between proteins in these complexes (47). A handful of studies have even identified that depletion of Zn^2+^ from the *E. coli* DNA polymerase I impairs its activity (48, 49); however, studying the role of these zinc finger domains in mammalian cells is prohibitively difficult. Since the subunits of the DNA polymerase complex are essential, and knockout or mutation of these genes in human cells is lethal, it is not possible to mutate these Zn^2+^ fingers in mammalian cells. That said, the evidence presented here rules out other causes of replication stress and suggests that DNA polymerase activity is sensitive to changes in the labile Zn^2+^ pool.

Replication stress in mother cells can be inherited by daughter cells to cause a pause in proliferation, accompanied by expression of p21 (4, 8, 35). Since impaired DNA synthesis, RPA2 foci accumulation, and Chk1 phosphorylation all decrease upon resupply of Zn^2+^, we sought to determine whether the memory of this DNA damage signaling influences the daughter cell’s proliferative fate. Cells exposed to a short pulse of TPA during the latter half of S-phase show an increase in cells entering a temporary CDK2 low state after cell division. These cells re-enter the cell cycle after a pause in proliferation, indicating that the inherited damage is reversible, and very few cells enter a permanent state of quiescence. This pause is accompanied by sustained expression of the tumor suppressor p21, which agrees with previous reports of inherited cell fate due to replication stress in the mother cell (4, 8, 35).

Using long-term live-cell imaging, fluorescent tools, and single-cell analyses, we identified a specific requirement for Zn^2+^ in S-phase. If cells are deficient in this essential micronutrient while synthesizing DNA, they show hallmarks of replication stress and DNA synthesis impairment. Further, this Zn^2+^ deficiency-induced replication stress impairs proliferation in daughter cells, maintained by sustained p21 expression. Together, these data demonstrate that mild and temporary Zn^2+^ deficiency can impair DNA synthesis within two hours and induce replication stress that is passed onto daughter cells after division.

## Materials and Methods

See Supplemental Information for detailed materials and methods.

### Cell culture

MCF10a cells (ATCC CRL-10317) were procured from ATCC and maintained in full growth media (FGM). FGM is composed of DMEM/F12 media supplemented with 5% horse serum, 1% penicillin/streptomycin solution, 20 ng/mL EGF, 0.5 μg/mL hydrocortisone, 100 ng/mL cholera toxin, and 10 μg/mL of insulin. Zn^2+^ defined minimal media (MM) consists of 1:1 Ham’s F12 nutrient/FluoroBrite DMEM, supplemented with 1.5% Chelex-100 treated horse serum, 10μg/mL Chelex-100 treated insulin, 1% penicillin/streptomycin solution, 20 ng/mL EGF, 0.5 μg/mL hydrocortisone, and 100 ng/mL cholera toxin. We generated MCF10a cells expressing PB-H2B-HaloTag (50) and the FUCCI (CA) (23) system using lentiviral transduction. MCF10a cells expressing H2B-mTq2, DHB-mCherry, and p21-mCitrine were provided by the laboratory of Dr. Sabrina Spencer at the University of Colorado Boulder, department of Biochemistry. See Supplemental Information for detailed materials and methods.

### Immunofluorescence

MCF10a cells were plated in 96-well plates at a density of 10,000 cells per well. Cells were treated and fixed with 4% paraformaldehyde in phosphate-buffered saline (PBS), and permeabilized with 0.1% Triton X-100 in PBS. Cells were then blocked in PBS with 3% bovine serum albumin (BSA) and stained with respective primary and secondary antibodies in PBS with 1% BSA. Images were collected via fluorescence microscopy. See Supplemental Information for detailed materials and methods.

### Image analysis

Time lapse image processing was done using EllipTrack (24). Data was analyzed with custom scripts. See Supplemental Information for detailed materials and methods.

## Acknowledgements

We would like to acknowledge NIGMS MIRA R35 GM139644 (A.E.P) for generous financial support. We thank the University of Colorado Flow Cytometry Core, which performed cell sorting and flow cytometric analysis of fixed cells, the University of Colorado Biochemistry Cell Culture Core (specifically Theresa Nahreini) for assistance with cell culture, and the BioFrontiers Computing Core and BioFrontiers IT for providing High Performance Computing resources. We also thank Joseph Dragavon for assistance with the microscopes available in the BioFrontiers Advanced Light Microscopy Core. We thank Dr. Luke Lavis (HHMI Janelia) for the JF669-HaloTag dye, and Dr. Sabrina Spencer (CU Boulder) for generously providing the MCF10a H2B-mTurquoise2, p21-mCitrine, DHB-mCherry cell line, and for the insightful comments on this manuscript. We also thank Dr. Shaun Bevers (University of Colorado School of Medicine Metabolomics Core, Aurora, CO) for preparing and analyzing the metabolomics samples.

