## Supplemental Information for "Transient Zn^2+^ deficiency induces replication stress and compromises daughter cell proliferation"

**This PDF file includes:**

Detailed Materials and Methods
Table S1
Figures S1 to S7
SI References

Supplemental Information

**Detailed Materials and Methods**

Table S1: Materials table

| Agent or Resource | Source | Identifier |
|---|---|---|
| Antibodies | | |
| Anti-RPA32/RPA2 antibody [9H8] | Abcam | ab2175 |
| Phospho Histone H3 (Ser10) (6G3) Mouse mAb | Cell Signaling Technology | 9706L |
| Anti-phospho Histone H2A.X Ser139 mouse monoclonal antibody | Cell Signaling Technology | 80312S |
| Anti-phospho Chk1 Ser317 rabbit | Abcam | ab226929 |
| Anti-phospho Chk2 Thr68 rabbit | Thermo Scientific | PA5-17818 |
| Purified mouse anti-human 53BP1 | BD Biosciences | 612523 |
| Mouse anti-MCM2 monoclonal antibody | BD Biosciences | 610700 |
| Goat anti-rabbit (H+L) Alexa Fluor 647 | Fisher Scientific | A21245 |
| Goat anti-mouse (H+L) Alexa Fluor 647 | Fisher Scientific | A21236 |
| Chemicals, peptides, and recombinant proteins | | |
| Tris(2-pyridylmethyl) amine 98% (TPA) | Sigma-Aldrich | 723134 |
| Chelex-100, sodium form | Sigma-Aldrich | C7901 |
| Janelia Fluor 669 Halo-Tag dye (JF669) | Laboratory of Luke Lavis, Janelia Research Campus | N/A |
| DMEM/F12, HEPES | Thermo Fisher Scientific | 11330057 |
| Horse Serum, New Zealand origin | Thermo Scientific | 16050122 |
| Hydrocortisone | Sigma-Aldrich | H4001 |
| Insulin | Life Technologies | 12585-014 |
| FluoroBrite DMEM | Fisher | A18967-01 |

| | | |
|---|---|---|
| Gibco EGF Recombinant Human Protein | Thermo Fisher Scientific | PHG0313 |
| Ham’s F12 phenol red-free | Sigma-Aldrich | N6658 |
| Cholera Toxin | Sigma-Aldrich | C8052 |
| Pen/Strep | Gibco | 15140-122 |
| 0.05% Trypsin-EDTA(1X) | Gibco | 25300-120 |
| Hoechst 33258 | Sigma-Aldrich | 861405 |
| FAM azide, 6-isomer | Lumiprobe | B5130 |
| 5-Ethynyl-2'-deoxyuridine,95% | Sigma-Aldrich | 900584-50MG |
| Alexa Fluor 647 azide | Lumiprobe | A6830 |
| L-Ascorbic acid | Fisher Scientific | A61-25 |
| Aphidicolin from Nigrospora sphaerica,>=98% (HPLC), powder | Sigma-Aldrich | A0781-1MG |
| Bleomycin, ready made solution | Sigma-Aldrich | B7216 |
| 4-Nitroquinoline N-oxide,≥98% | Sigma-Aldrich | N8141-250MG |
| CHIR-124 2mg | Selleck Chemicals | S2683 |
| Critical Commercial Reagents | | |
| TransIT-LT1 | Mirus Bio | MIR 2300 |
| Experimental models: cell lines | | |
| Human: MCF10a | ATCC | CRL-10317 |
| Human: MCF10a + H2B-HaloTag | Rakshit and Holtzen 2023 (1) | PMID: 37330912 |
| Human: MCF10a + H2B-HaloTag + FUCCI (CA) | This work | N/A |
| Human: MCF10a + H2B-mTq2 + DHB-mVenus | Lo 2020 (2) | PMID: 32014109 |
| Human: MCF10a + H2B-mTq2 + DHB-mCherry + p21-mCitrine | Moser 2018 (3) | PMID: 30111539 |
| Recombinant DNA | | |

| | | |
|---|---|---|
| Plasmid: PB-H2B-HaloTag (used for stable cell line generation) | Grimm 2017 (4) | PMID: 28869757 |
| mVenus-hGeminin(1/110)/pCSII-EF | Riken | RDB15271 |
| mCherry-hCdt1(1/100)/pCSII-EF | Riken | RDB15442 |
| Software and Algorithms | | |
| BD FACSDiva Version 8 | BD Biosciences | N/A |
| MATLAB 2017a and VR2020b | Mathworks | N/A |
| Prism Version 9.2.0 | GraphPad | N/A |
| EllipTrack | Tian 2020 (5) | PMID: 32755578 |

Cell culture

MCF10a cells were procured from ATCC and maintained in full growth media (FGM). FGM is composed of DMEM/F12 media supplemented with 5% horse serum, 1% penicillin/streptomycin solution, 20 ng/mL EGF, 0.5 µg/mL hydrocortisone, 100 ng/mL cholera toxin, and 10 µg/mL of insulin. Cells were passaged at 80-90% confluency using 0.05% trypsin-EDTA. To remove excess $Zn^{2+}$ from horse serum and insulin, we incubated serum and insulin solutions with Chelex-100 resin for 12 hours at 4ºC, followed by sterile filtration to remove resin. Our $Zn^{2+}$ defined minimal media (MM) consists of 1:1 Ham's F12 nutrient mix:FluoroBrite DMEM, supplemented with 1.5% Chelex-100 treated horse serum, 10µg/mL Chelex-100 treated insulin, 1% penicillin/streptomycin solution, 20 ng/mL EGF, 0.5 µg/mL hydrocortisone, and 100 ng/mL cholera toxin. We previously quantified metal content of MM containing Chelex-100 treated serum, which showed that Chelex-treatment leads to a reduction in zinc (from 2.18 to 1.46 µM) and nickel (from 0.178 to 0.012 µM), but no significant changes in other metals (2). Unless otherwise stated, ZD media was generated by adding 3 µM TPA to MM. For imaging experiments, cells were grown and imaged in MM, with perturbations with ZD media at 37ºC and 5% $CO_2$. Cells used in experiments were less than passage number 15 and were routinely tested and confirmed to be mycoplasma negative by PCR.

Plasmids and cell lines

MCF10a cells expressing PB-H2B-HaloTag (4) and the FUCCI (CA) (6) system were generated using lentiviral transduction. Briefly, HEK293T cells were transiently transfected with both the hGem-mVenus and hCdt1-mCherry plasmids, along with Lenti-X lentiviral packaging plasmids psPax2, PdM2.G in OptiMEM using the TransIT-LT1 transfection reagent (Mirus Bio). This was followed by viral amplification in HEK293T cells Successful transfection was verified via visualization of mCherry and mVenus fluorescence in the HEK293T cells. Viral particles were harvested 24 hours after transfection, filtered through 0.45 µm syringe polyethersulfone (PES) filters and then 1 mL was added to MCF10a cells stably expressing H2B-HaloTag and allowed to incubate for 48 hours along with 4 µg/mL polybrene. Stable cell lines used for imaging were then FACS sorted on a BD FACSAria Fusion for high mCherry and mVenus fluorescence relative to a non-transduced control using the following optics: YFP: Ex 488 nm, Em 530/30 nm; and mCherry Ex 561 nm, Em 610/20 nm.

Live-cell imaging

MCF10a cells expressing fluorescent reporters were counted using the Countess II Automated Cell Counter (Thermo Fisher Scientific) and plated in MM into glass bottom 96-well plates (Cellvis P96-1.5H-N) at a density as to ensure cells were subconfluent by the end of the time lapse. After plating, cells were left undisturbed in the biosafety cabinet for 30 minutes to properly adhere to the dish, then moved to the incubator to recover for at least 16 hours. Immediately before imaging, cells expressing H2B-HaloTag were stained with 5 nM JF669 dye (Luke Lavis, Janelia Research Campus) for 30 minutes, then washed once with prewarmed MM. For the ZD pulse-chase experiment, acquisition was paused, half of the media was removed, and replaced with a 2X solution of prewarmed ZD media. To wash out the treatment, 90% of the media was removed from the well and replaced with prewarmed MM. For the “To End” condition, treatment media was not removed.

Time-lapse images were collected using a Nikon Ti-E High Content Analysis inverted microscope with a Lumencor SPECTRA X light engine (Lumencor) and Hamamatsu Orca FLASH-4.0 V2 scMOS camera (Hamamatsu). Images were collected every 12 minutes with a 10X 0.45 NA Plan Apo air objective lens (Nikon Instruments). During imaging, cells were kept in a controlled environmental chamber surrounding the microscope (Okolab Cage Incubator, Okolab) at 37ºC and 5% $CO_2$ and 90% humidity. Filter sets for live-cell imaging were as follows: CFP Ex: 440, 455 dichroic, Em: 480/20, power 50; YFP Ex: 508, 518 dichroic, Em: 540/21; mCherry Ex: 555, 597 dichroic, Em: 595/40; and JF669/Cy5 Ex: 640, 640 dichroic, Em: 705/22, power 50. Exposure times were adjusted to eliminate pixel saturation and to minimize exposure time, while still maintaining an adequate signal to noise ratio.

Time-lapse image processing

Live-cell imaging experiments were analyzed using the EllipTrack cell tracking pipeline in MATLAB 2017a and 2020b on the institutional computing cluster (5). Briefly, EllipTrack requires several generalized parameters for cell segmentation, tracking and event identification. The advanced parameters were unchanged from the original code accessible on Github, as recommended by the authors. Basic parameters were fine-tuned by hand using the built-in user interface. Separate training data sets for the EllipTrack event predictions were made using at least 500 events across each time series for both H2B-HaloTag and H2B-mTurquoise2 nuclear markers. We used a nuclear radius of 12 pixels as the average size of one cell. The ellipse that is fitted to this nucleus was required to be at least 25 pixels in area. Nuclear images were log transformed and a blob detection algorithm was used to identify nuclei with a blob threshold of -0.075. Objects were separated using a watershed algorithm and ellipses were fitted to each object. Mitoses were inferred based on morphological properties of each object before and after a potential mitosis event. Migration and migration speed were inferred using a density-dependent migration speed from the training data. All tracks greater than 10 frames were kept, and tracks were allowed to skip at most two frames. Tracking was spot-checked by eye using the visualize tracking (vistrack) movies generated by the script to ensure adequate tracking and minimize track loss or swapping. Mean intensity from nuclear localized reporters was extracted using the nuclear mask. The mean intensity of cytoplasmic sensors was calculated in a three-pixel wide cytosolic ring around the nuclear mask.

Flow Cytometry EdU Incorporation Assay

Cells were plated at sub-confluency in 6-well plates in MM and allowed to recover for at least 16 hours. Cells were treated with ZD media for 2-4 hours, co-treated with ZD media and 250 or 500 nM CHIR-124, or left untreated. Thirty minutes before harvesting, cells were pulse-labeled with 10 µM EdU for 30 minutes in the cell culture incubator. Cells were then harvested using 0.005% EDTA/Trypsin and pelleted at 300 x g for 5 minutes. Cells were fixed using 4% paraformaldehyde (PFA) and washed twice with phosphate buffered saline (PBS) + 1% bovine serum albumin (BSA) (B-PBS). Cells were permeabilized with 0.2% Triton-X100 for 15 minutes, then washed twice with B-PBS. Cells were then resuspended in a click solution consisting of 50µM FAM-Azide, 2 mM $CuSO_4$, and 50 mM L-ascorbic acid in PBS for 30 minutes in the dark. Cells were washed with B-PBS and stained with an analysis buffer (B-PBS with 1:10000 dilution of 1 mg/mL DAPI) for 1h. Cells were analyzed on a BD FACSCelesta outfitted with 405 and 488 nm optics. FACS files were exported and analysis was done in FlowJo or at floreada.io, a free open-source flow cytometry analysis tool. S-phase was identified by gating EdU+ cells. The median EdU signal was extracted from this population in all treatment conditions. These signals were then normalized to the median EdU signal of the untreated control.

Loaded MCM2 Assay

Fixed cell immunostaining of loaded MCM2 and flow cytometric analyses were performed as previously described (7). Briefly, following treatments or release from serum starvation, cell suspensions were fixed with 4% PFA and washed twice before stepwise staining: EdU click reaction with FAM-Azide (room temperature for 30 minutes), mouse anti-MCM2 immunostaining (BD Biosciences #610700, 1:200 dilution in PBS with 1% BSA for 1 hour at 37ºC), goat anti-mouse Alexa Fluor 594 immunostaining (Thermo Fisher A-11005 for 1h at 37ºC), and Hoechst 33342 (1:10,000 in PBS overnight at 4ºC). Cell suspensions were filtered for single cells and analyzed using a BD FACSCelesta flow cytometer outfitted with 405, 488, and 561 nm lasers. FACS files were exported and analyzed using FlowJo or at floreada.io, a free open-source flow cytometry analysis tool.

Cells were first gated for cells on forward scatter area vs. side scatter area, then gated on DAPI signal height vs. DAPI signal area for single cells. MCM2+ S-phase cells were then identified using plots of EdU incorporation vs loaded MCM2 in the untreated control. These gates were then applied to all other cells to identify the fraction of cells that entered early G1 with under-licensed origins of replication.

Detection of mitotic DNA synthesis (MiDAS)

MCF10a cells were plated in 96-well plates at a density of 10,000 cells per well into MM. Plates were left in the biosafety cabinet for 30 minutes to allow cells to adhere to the dish, then transferred to the incubator. After 4 hours, positive control cells were treated with 0.2 μM aphidicolin for 24 hours. Cells were treated with 3 μM TPA for 4 or 6 hours. For the resupply experiment, ZD media was removed from cells and replaced with fresh MM. Media was then removed from all cells and replaced with media containing 40 μM EdU, and allowed to incubate for 30 minutes. Cells were washed with PBS and fixed with 4% PFA in PBS for 10 minutes, then permeabilized with PBS containing 0.1% Triton-X100 for 10 minutes. Cells were washed and incubated with a click chemistry buffer containing Alexa Fluor 647 azide as described previously. Cells were then blocked using PBS containing 3% BSA for 1h before staining with mouse anti-phospho histone H3 (Ser10) (1:400 dilution into PBS + 1% BSA, Cell Signaling Technologies) overnight at 4ºC. Cells were then washed and counterstained with a goat anti-mouse secondary antibody conjugated to Alexa Fluor 488 (1:1000 into PBS + 1% BSA, Thermo Fisher) and DAPI. Cells were washed and then immediately imaged as described below.

Immunofluorescence and Fixed Cell Imaging

MCF10a cells expressing H2B-HaloTag and the FUCCI (CA) system were plated in 96-well plates (for Chk1 IF and yH2AX IF, cells were plated in Cellvis P96-1.5H-N plates, for RPA2 IF, cells were plated in PerkinElmer PhenoPlate 96) at a density of 10,000 cells per well. Plates were left in the biosafety cabinet for 30 minutes to allow for proper adhering, then allowed to recover in the incubator for at least 16 hours undisturbed. Prior to fixation, cells were treated for either 2h or 4h with ZD media, or 4 hours with 2 μg/mL aphidicolin. For EdU incorporation at short timepoints, cells were pulsed with 10 μM EdU 30 minutes before fixation. At the time of fixation, cells were washed twice with PBS pH 7.4, then fixed with 4% PFA in PBS pH 7.4 for 15 minutes. Cells were then washed twice with PBS, then permeabilized with 0.05% Triton-X100 in PBS for 20 minutes. Cells were then washed twice with PBS and blocked for 1 hour with PBS and 3% bovine serum albumin (BSA). Afterwards, the BSA solution was removed and the primary antibody solution (1:250 for all targets in 3% PBS) was added without washing. This was allowed to incubate overnight for at least 16 hours at 4ºC. Cells were washed 4X with PBS pH 7.4, then stained with secondary antibody (1:1000 into PBS + 1% BSA goat anti-mouse (H+L) Alexa Fluor 647 for RPA2 and yH2AX, and 1:1000 goat anti-rabbit (H+L) Alexa Fluor 647 for Chk1 pSer317) in PBS with 3% BSA and 100 ng/mL Hoechst dye for 1h at room temperature. Cells labeled with EdU alone were stained using click chemistry to attach an Alexa Fluor 647 azide dye to the ethynyl handle of EdU as described previously. Cells were immediately imaged.

Images of pChk1, yH2AX, and EdU were collected using a Nikon Ti-E High Content Analysis inverted microscope with a Lumencor SPECTRA X light engine (Lumencor) and Hamamatsu Orca FLASH-4.0 V2 scMOS camera (Hamamatsu) using a 40X 0.65 NA CFI Plan air objective from Nikon. A 9x9 large image was taken of the center of each well of the 96-well plate. The DAPI channel was acquired using a 395/15 excitation / 475/24 emission. The YFP channel was acquired using a 510/25 excitation 540/21 emission filter set. The mCherry channel was acquired using a 575/25 excitation 632/60 emission filter set. The Cy5 channel was acquired using a 640/30 excitation 705/72 emission filter set. Single-cell quantitation was done by identifying nuclei using a custom MATLAB pipeline. Since these targets are localized to the nucleus, we then extracted mean intensity in the DAPI, YFP, mCherry, and Cy5 channels in the region of the identified nuclear masks.

Images of RPA2 were collected on a PerkinElmer Opera Phenix high-content screening system with a 20x/1.0 NA water objective. Eight images of each well were collected, each containing three Z-stack images of all channels. Maximal intensity projections were calculated, and used in subsequent analysis of all channels. Single-cell quantitation was done by identifying nuclei using a custom MATLAB pipeline, then extracting mean intensity in the DAPI, YFP, and mCherry channels in those regions. Additionally, single RPA2 foci were found by adaptive thresholding of each nuclear object on the Cy5 channel, with area filtering of objects greater than 5 pixels and with an eccentricity of less than 0.8.

Images of MiDAS were collected on a PerkinElmer Opera Phenix high-content screening system using the PreciScan modality. Briefly, cells were imaged at a low magnification (10X/0.4 NA air objective), then cells were identified and selected for high phospho-histone H3 (Ser10) fluorescence. These cell locations were recorded and the objective was switched to a higher magnification (40X/1.1 NA water objective) and imaged. Six Z planes were acquired for each cell, and at least 20 regions were imaged per well, yielding at least 150 cells per condition. Image processing and MiDAS foci selection was done using the PerkinElmer Harmony software.

Relative metabolite quantification

Relative metabolite quantification was performed by the University of Colorado School of Medicine Metabolomics Core (Aurora, CO). Metabolite extraction was accomplished by the addition of 4°C lysis buffer (LB) (5:3:2 methanol:acetonitrile:water (v/v/v)) to a concentration of $2 \times 10^6$ cells/mL (based on approximate cell counts) to frozen cell pellets. This mixture was vortexed for 30 min at 4°C and centrifuged for 10 min (18,000g, 4°C) to re-pellet the cells. The supernatant was then removed and placed in autosampler vials for UHPLC-MS analysis. A Thermo Vanquish UHPLC system combined with a Thermo Orbitrap Exploris 120 mass spectrometer (Thermo Fisher Scientific, Waltham, MA, USA) was used for untargeted metabolite analysis. Samples were analyzed both in positive and negative ion mode in separate runs using a 5 min gradient method (8) (7).

El-Maven v0.12.0 was used to identify, quantitate (by peak area) and annotate mass spectra features to produce a global metabolomic analysis (9). Statistical analysis (PLS-DA, Heatmap) and pairwise comparisons were accomplished using MetaboAnalyst 5.0 (10).

Data analysis

*Analysis of the DHB-mVenus pulse-chase experiment:*

Single cell tracks were extracted from the EllipTrack output, and each mother and daughter cell was joined by at least one mitosis event to get combined mother/daughter cell tracks. To remove non-expressing cells and tracks that were swapped during analysis, two filters were employed: in the first, cells were removed that had an intensity value below a cutoff chosen for each replicate of the experiment. Second, cells that were swapped during tracking have at least one very short intermitotic time (IMT), indicating that the track was attributed a mitosis when there was not one, or the track was swapped with a cell that was mitotic. An IMT cutoff of 50 frames was used, since MCF10a cells have a cell cycle of 60 frames or longer. A sampling of tracks were analyzed by eye to ensure the analyzed tracks behaved as expected, and that poorly tracked cells were kept to a minimum.

CDK2 activity was calculated at every frame by dividing the mean cytoplasmic signal of the DHB-mVenus reporter by the mean nuclear signal. This will typically yield a CDK2 activity ranging between 0.2 and 2 AU. To identify the cell cycle dependence of $Zn^{2+}$ in cells, tracks were binned based on the frame at which they divided. For each frame, we calculated the cell fate of cells dividing in a 5 frame window around that frame in the following manner: if cells had a CDK2 activity > 0.5 by 15 frames after mitosis, they were called "$CDK2_{inc}$". If cells had a CDK2 activity < 0.5 15 frames after mitosis, they were either $CDK2_{emerge}$ or $CDK2_{low}$. To delineate, we determined whether cells began building up CDK2 activity after division, or if they stayed low for the rest of the time lapse. The average and 95% confidence interval was determined for each frame. The data were then plotted as "drug addition relative to anaphase". Since proliferating cells have an IMT of between 60-70 frames, or 12-14 hours, the time point -12 is taken to be either mid or late G1. After cells commit to a cell cycle, the time from S-phase to mitosis is very robustly 8 hours. All analysis was conducted in MATLAB 2020B.

*Analysis of the DHB-mCherry p21-mCitrine pulse-chase experiment:*

Single-cell tracks were constructed identically to the DHB-mVenus pulse-chase experiment. Mean nuclear fluorescence of the p21-mCitrine signal was used to determine p21 protein expression at each frame of the time-lapse. For each treatment condition, cells were binned according to their CDK2 activity after division similarly to the DHB-mVenus pulse-chase experiment. Since cells were expressing DHB-mCherry, the threshold for $CDK2_{inc}$ and $CDK2_{emerge}$ cells was 0.58 instead of 0.5, which we identified empirically. The CDK2 activity and p21 signal was then averaged and 95% confidence intervals were calculated.

*Analysis of the FUCCI (CA) pulse-chase experiment:*

Tracks of cells expressing the FUCCI (CA) sensor had to exist for the entirety of the timelapse and were joined by every mitosis event detected by the EllipTrack program. Cells not expressing both FPs at some point in the time lapse were removed from analysis. To identify mitosis events, the local minima were found in the derivative of the mVenus signal. To identify cell cycle phases, the mVenus and mCherry signals were min-max normalized, and subjected to hysteresis thresholding to find roughly where each sensor was on or off. These thresholded FUCCI (CA) signals were then passed through a first-round phase calling algorithm, which simply assigned the G1 phase to mCherry(+) cells, S-phase to mVenus(+) cells, and G2 to mCherry(+) and mVenus(+) cells. Occasionally, this algorithm miscalled the G1/S transition as G2, so a refining function was applied to ensure that cells progressed from G1 to S to G2 to M to G1 and so on in that exact order. Cell cycle phase lengths were measured for each mother and daughter cell and exported for analysis. All analysis was done in MATLAB 2020B and in Python.

*Identifying FUCCI cell cycle phase using a Gaussian Mixture Model (GMM)*

Mean nuclear intensities of the FUCCI reporter were extracted from fixed cells in every IF experiment. The z-score of each intensity was calculated for each replicate, and then each cell was categorized using a GMM with a set random seed and 10,000 iterations, which ensured that the model converged. Cells were then plotted and color-coded with their respective predicted group, and cell cycle phase was assigned by eye for each experiment. Cells in the group of mCherry(-) and mVenus(-) were removed from analysis. Analysis was done in Python.

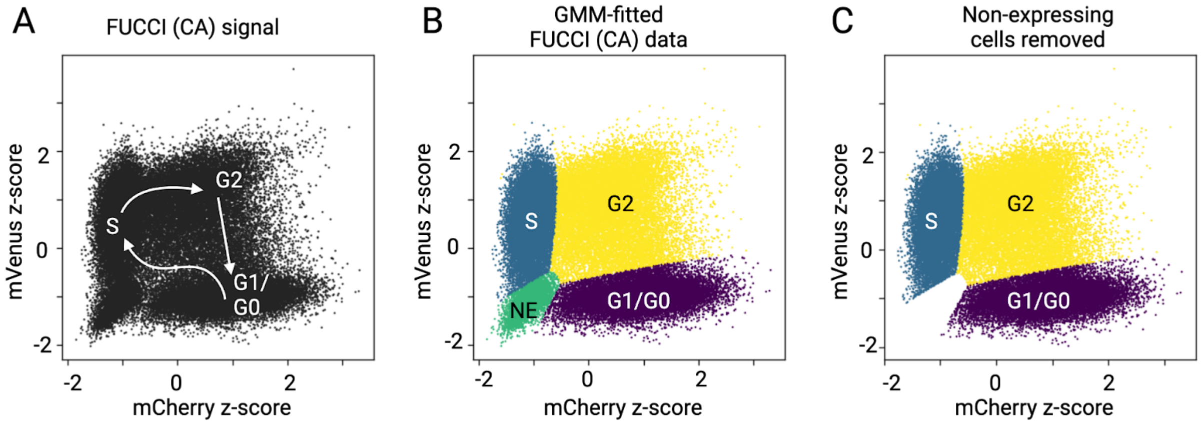


**Figure S1: A Gaussian Mixture Model identifies cell cycle phases in fixed cells expressing the FUCCI (CA) sensor.** (A) Quantitative imaging-based cytometry plots showing the z-score of mean nuclear fluorescence of single cells expressing the FUCCI (CA) sensor. Cells with low mVenus intensity and high mCherry intensity are in G1/G0. Cells with high mVenus intensity and low mCherry intensity are in S-phase. Cells with both high mVenus and mCherry intensities are in G2. Arrows indicate the trajectory of cells throughout the cell cycle in pseudotime. (B) A Gaussian mixture model was fitted to the data in A and cell cycle phases were color coded and assigned by hand. (C) Cells with low mVenus and low mCherry intensities were removed from further analysis.

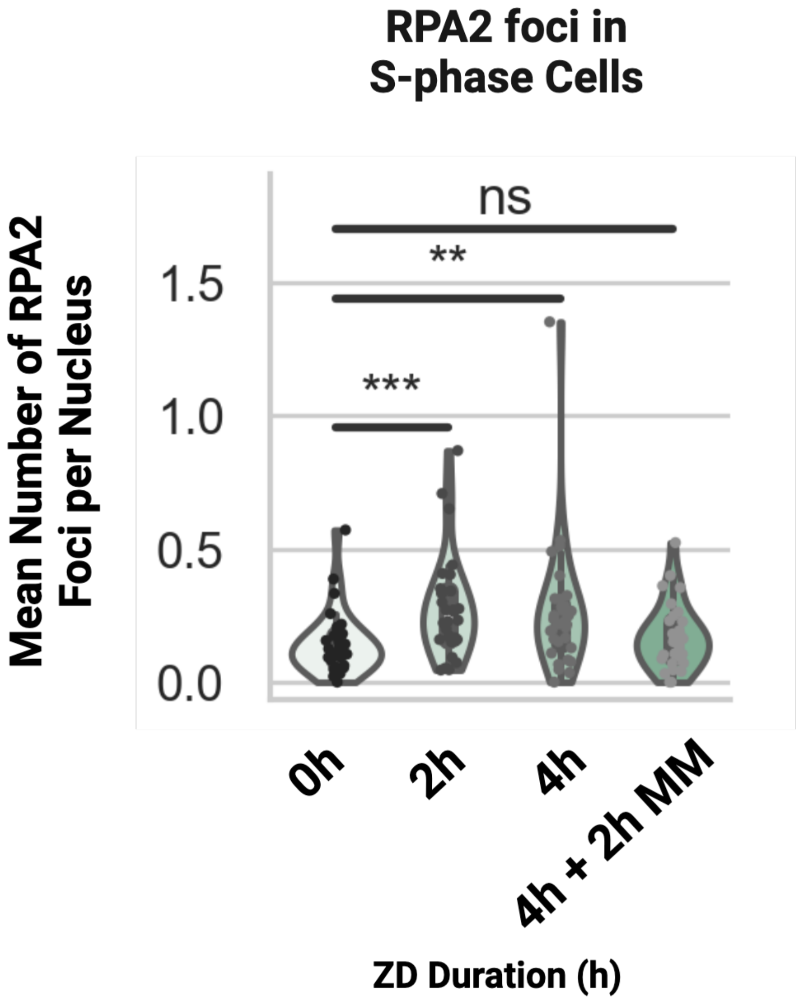


**Figure S2: RPA2 foci in $Zn^{2+}$ deficient cells resolve after resupply of $Zn^{2+}$.** Violin plots of mean number of RPA2 foci per nucleus for cells in S-phase as determined by immunofluorescence. $Zn^{2+}$ deficient duration on the horizontal axis. *** = $p < 0.005$, ** = $p < 0.01$, ns = not significant. n = 3 biological replicates.

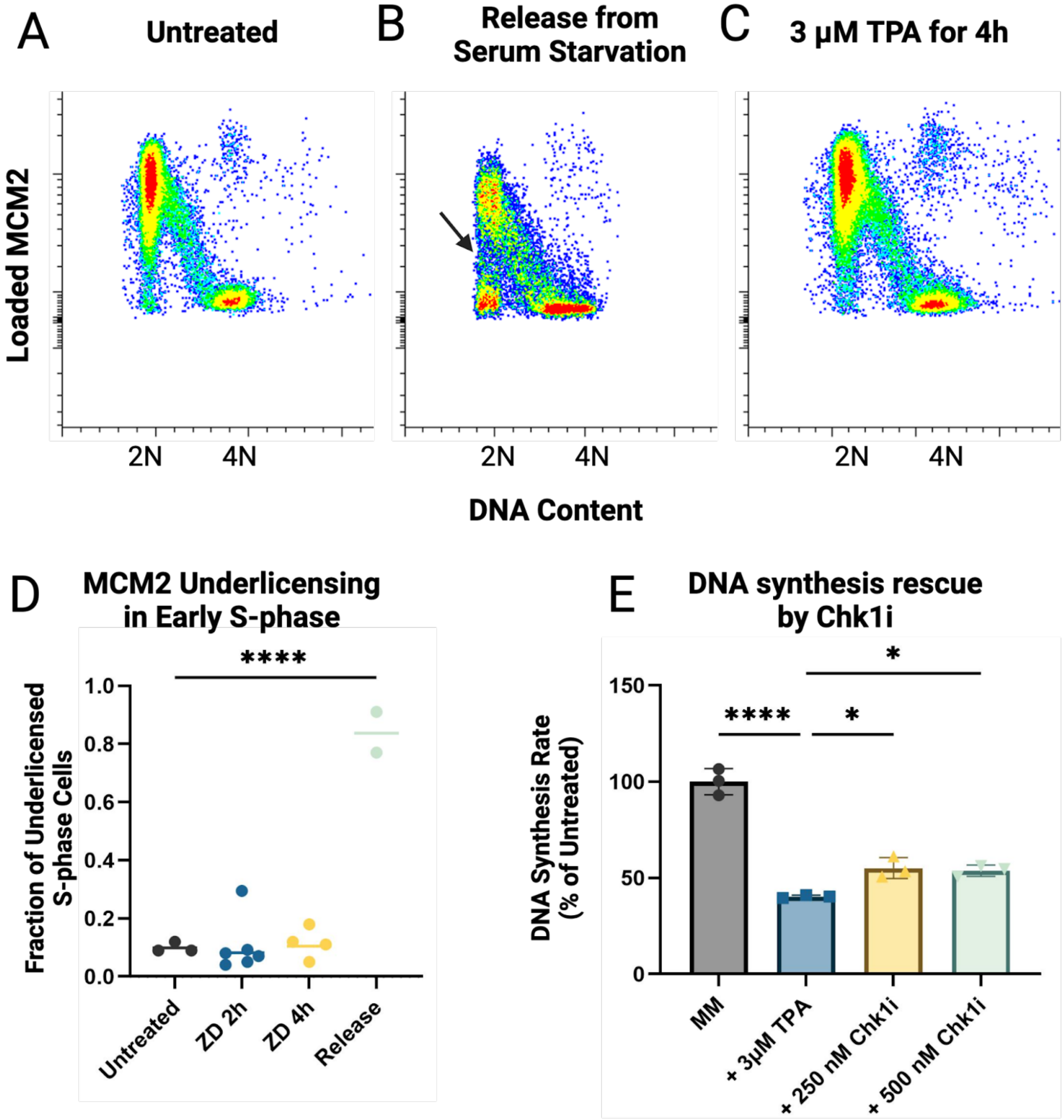


**Figure S3: $Zn^{2+}$ induced DNA synthesis impairment is not due to under licensing of origins of replication, nor is it fully due to origin firing inhibition by Chk1.** (A) Flow cytometry plots of untreated cells showing chromatin loaded MCM2 and DNA content using single nuclei stained with an anti-MCM2 antibody and DAPI. Graphs show characteristic increase in MCM2 loading during G1, and decreased MCM2 loading throughout S-phase. (B) Flow cytometry plots of cells released from serum starvation showing chromatin loaded MCM2 and DNA content. Cells released from serum starvation show under licensing of replication origins, as shown with the arrow, and as shown previously. (C) Flow cytometry plots of $Zn^{2+}$ deficient cells showing chromatin loaded MCM2 and DNA content. (D) Fraction of under-licensed S-phase cells in each condition using a previously established analysis scheme (7). (E) Quantification of DNA synthesis rate like that shown in figure 2B. Cells were treated with 3μM TPA for 4h, or 3μM TPA and the Chk1 inhibitor CHIR-124, then probed for DNA synthesis rate using EdU. Each point is the median EdU signal of S-phase cells, normalized to untreated control. Error bars represent the 95% confidence interval. * = $p<0.05$, **** = $p < 0.001$. $n \geq 2$

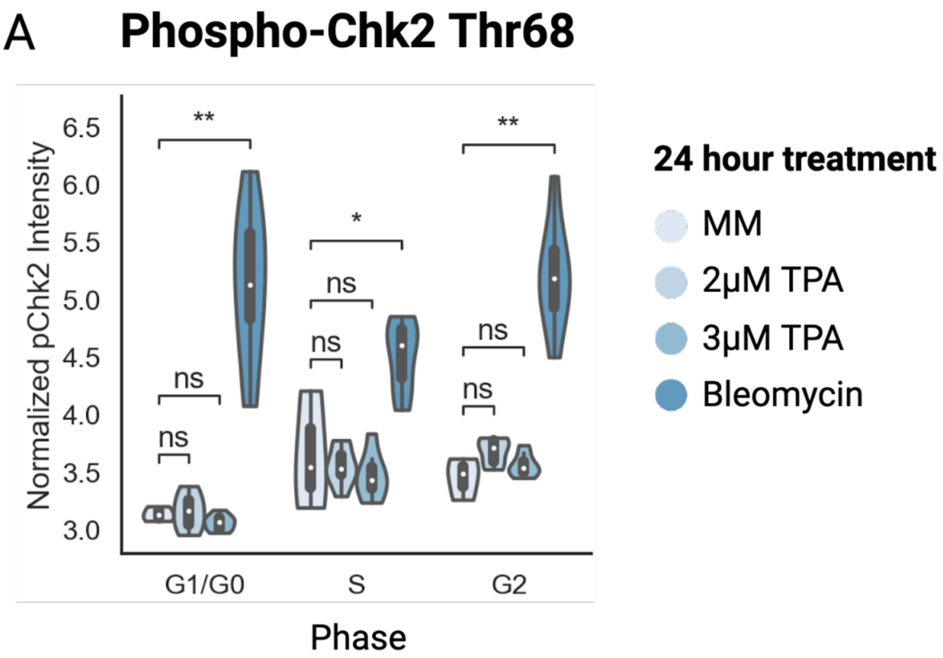


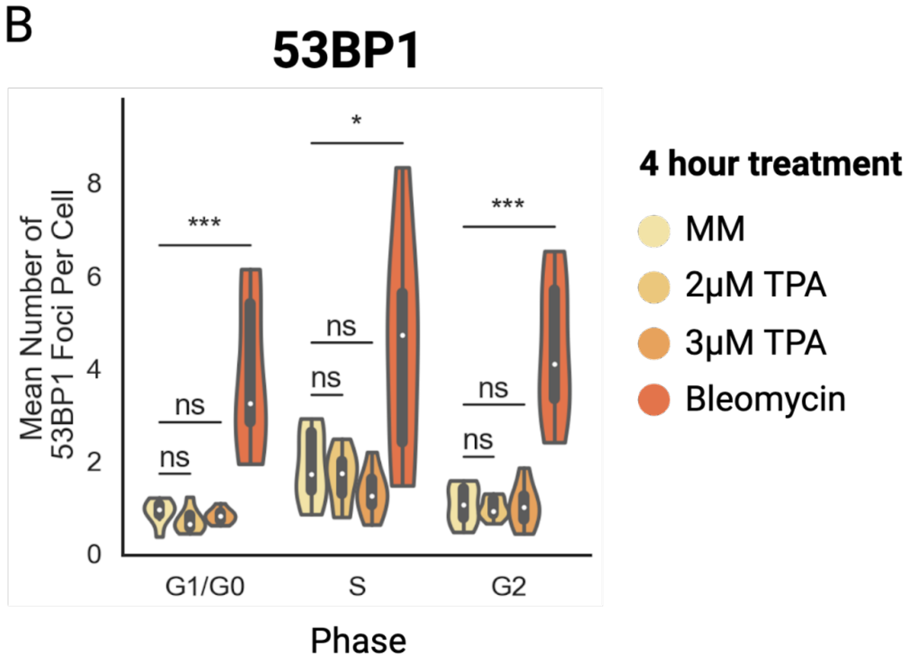


**Figure S4: $Zn^{2+}$ deficiency does not activate Chk2 nor does it induce double-stranded breaks** (A) Quantification of 53BP1 foci in each cell cycle phase after four-hour treatment with $Zn^{2+}$ deficient media. Each data point is the average number of 53BP1 foci in cells in that phase in one well of a 96-well plate. Bleomycin 4μM is used as a positive control. (B) Quantification of phospho-Chk2 (Thr68) signal in S-phase cells after 24-hour treatment with $Zn^{2+}$ deficient media. Each data point is the average log transformed nuclear Chk1 phospho-Thr68 in cells in that phase in one well of a 96-well plate. Bleomycin 4μM is used as a positive control. * = $p < 0.05$, ** = $p < 0.01$, *** $p < 0.005$. n = 3 replicates. Cell cycle phases identified using nuclear FUCCI (CA) signal and a Gaussian mixture model (GMM).

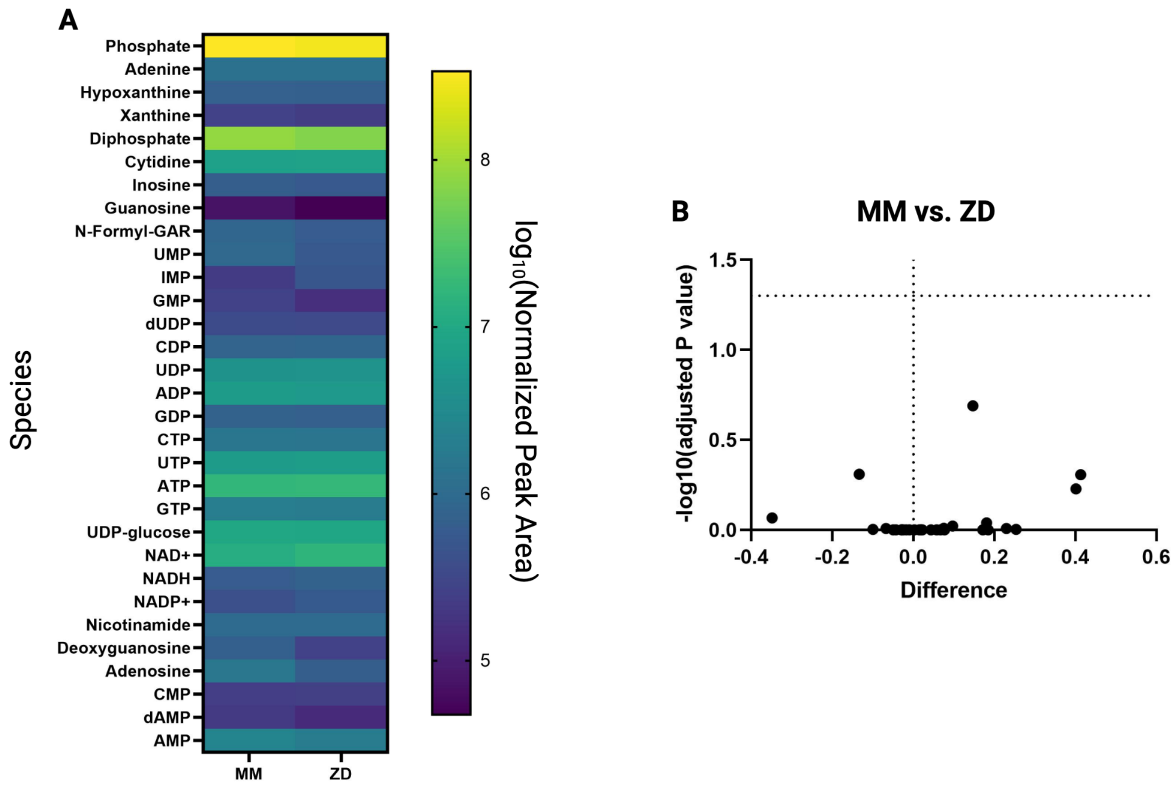


**Figure S5: $Zn^{2+}$ does not cause imbalance or depletion of nucleotides or their precursors.** (A) Heatmap of the average $\log_{10}$ peak area of each metabolite involved in the nucleotide biosynthesis pathway. MM indicates untreated cells, and ZD indicates cells treated with 3μM TPA for four hours. (B) A volcano plot showing changes in relative abundance of each species, and the adjusted P value for each. The P-values are adjusted using the Benjamini-Hochberg multiple hypothesis correction. The horizontal dotted line indicates threshold for significance. No species exceeds the adjusted P-value threshold. Each condition contains n=5 biological replicates.

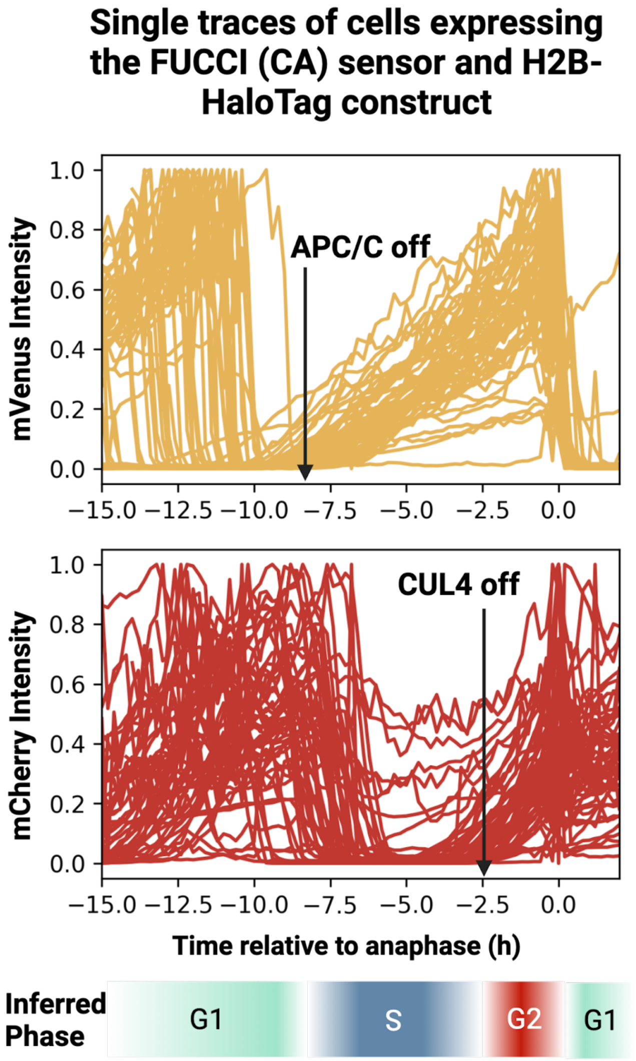


**Figure S6 The FUCCI (CA) sensor can identify cell cycle phase transitions at a population level.** (Top) Single-cell traces of the mVenus component of the FUCCI (CA) sensor aligned to mitosis. The black arrow indicates the time at which the population of cells enters S-phase, as evidenced by the buildup of the mVenus signal. (Middle) Single-cell traces of the mCherry component of the FUCCI (CA) sensor aligned to mitosis. The black arrow indicates the time at which the population of cells enters G2, as evidenced by the buildup of the mCherry signal. (Bottom) The inferred phase of the cell cycle as a function of time relative to mitosis.

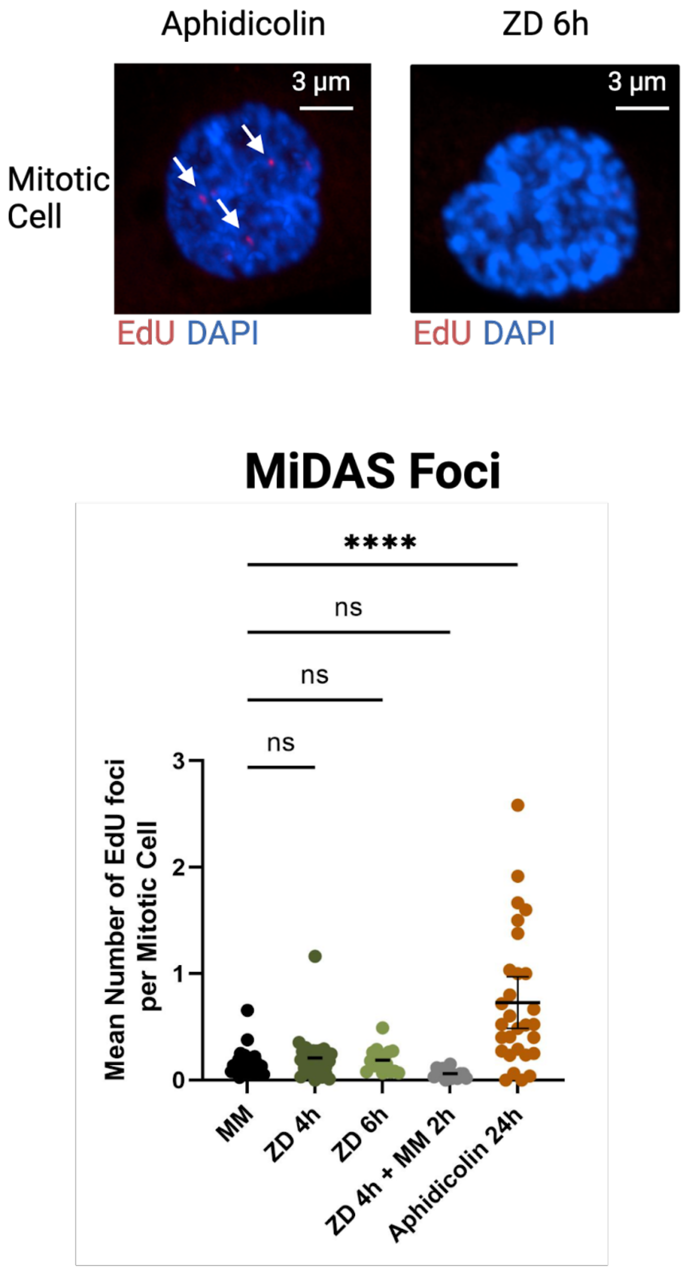


**Figure S7 Neither $Zn^{2+}$ deficiency nor $Zn^{2+}$ resupply stimulates mitotic DNA synthesis (MiDAS).** (Top) Representative images of mitotic cells treated with either 0.2 µM aphidicolin for 24h or $Zn^{2+}$ deficiency for 6 hours. Note the red EdU foci in the aphidicolin treatment (white arrows) and absence thereof in the $Zn^{2+}$ deficient treatment. (Bottom) Quantification of MiDAS foci. Each dot represents average MiDAS foci in one well of a 96-well plate. Error bars represent the 95% confidence interval. **** = $p < 0.001$, $n \geq 2$ biological replicates.
